# Bistable host-pathogen interaction explains varied infection outcomes

**DOI:** 10.1101/2021.04.13.439629

**Authors:** Guozhi Yu, Yingying Hu, Shizhong Wang, Xingfa Han, Xiaogang Du, Huailiang Xu, Xianyin Zeng, Ulrich Steiner, Jens Rolff

## Abstract

Death or survival are the two outcomes of an infection. The development of an infection depends on multiple interplaying factors, including the temporal dynamic of host nutrition, immune response and pathogen proliferation, but how these factors collectively determine the outcome of an infection is less understood. To understand the underlying principle of infection, here, we took a systems approach by developing a simple mathematical model to understand multi-factored host-pathogen interaction. We find that nonlinear interactions feedback between host and pathogens resulting in two distinct trajectories of disease progression. We show that disease progression, measured by the pathogen load, responds in a bistable manner, by either proliferating or eventually being eliminated. We identified key processes accounting for bistability, including host’s resource assimilation, immune response and intrinsic noise. In addition, resource availability, immunity, and pathogen load at the onset of infection are also critical in determining the bistability. We then highlight, by discussing in the light of previous theoretical and experimental work, how our framework provides testable theories for host-pathogen interaction inside the host.

## INTRODUCTION

Infected hosts either die or survive. The progression of an infection and its symptoms depend on various factors that have been qualitatively examined in numerous ecological immunology studies [1–5]. For example, manipulating nutrition and resource availability mediates host resistance and tolerance to pathogen infection, and thereby determines the course of infection [3,6–8]. Hosts consuming protein-rich diet resist bacterial infection more efficiently and survive better than hosts on a protein-deprived diet [9,10]. The protein-rich diet enhances the production of immune effectors such as antibodies [11] and antimicrobial peptides [5]. Carbohydrate-rich food by contrast can reduce the host’s ability to eradicate pathogens but may result in hosts harboring more pathogens before the onset of symptoms [5], but see [13]. Experiments on diet-restriction, diet choice and metabolic shift during infection have also highlighted the significant role of resource and nutrition in host immunity [6,14–17]. In addition, the genetic background of both host and pathogen can affect the outcome of infections even within the same or closely related species [18]. For instance, hosts with varied and diversified expression of antimicrobial peptides differ in their ability in either resisting or tolerating pathogens [19–21].

To disentangle the contribution of multiple factors on the outcome of infection, a common approach is to analyze experimental studies that apply a full-factorial design by general linear models. However, such analyses have been criticized as they are limited to linear treatment-response hypotheses, single time point sampling, and their difficulties in inferring causality [22–24]. Within-host pathogen dynamics tend to be nonlinear, which means response variables do not proportionately change with explanatory factors or time [25,26]. For example, the progression of malaria infections in mice revealed damped oscillations of both pathogen load and blood immune cell count [27]. In other words, both the pathogen population and host’s health fluctuate during the course of an infection. Measuring multiple response variables of the same host at different time points during infection clearly revealed such temporal dynamics [34]. Another important form of non-linearity is bistability, for which the pathogen load and host health are eventually attracted to two “steady states” corresponding to the death and survival of the host, respectively [25]. Empirical data, combined with mathematical analyses, revealed that the progression of infection and health dynamics in a number of hosts show this pattern [25,28–30]. However, in lab experiments, such within-host temporal dynamics can be masked in small animal models that cannot be sampled more than once, for instance when sampling is destructive. Pooled sampling, by sacrificing enough individuals at each time point, still revealed bistability in pathogen load in *Drosophila melanogaster* [31]. In experiments with a defined inoculum, pathogens firstly propagate, then either reach the carrying capacity of the host, or decrease to very low level. This divergence beginning at some time point can be shown with time series data [31].

Therefore, a theoretical and practical framework of host-pathogen interactions and feedbacks could provide an understanding of the within-host infection dynamics [28,32,33]. By including the concept of within-host resource competition in host-pathogens dynamics, researchers have built models that capture the distribution of host resources in an infection [32,34]. Possible feedbacks between resource, host immunity, and pathogens are likely to be the vital factors affecting the dynamics. To gain a quantitative understanding of within-host dynamic of infection, we extend the within-host resource-centric framework to host-pathogen interactions and their feedbacks. Our framework integrates the type of infection, and the regulation of the pathogen load inside the host via host resources. Our model recapitulates that host fitness is reduced when the pathogen load is high, i.e. causing sickness behaviors or changing food consumption. The host immune system is activated by pathogen invasion, and immediately targets and finally can eliminate the pathogens, mediated by metabolic adjustment. Most importantly, the energetic and resource costs of the dynamics are balanced by the host’s ingestion and metabolism. Thus, we expect a tipping point in the dynamics that differentiates the sustainability and collapse of the system, which also determines the survival or death of the host. In order to explain the phenomenon, we build a simple mathematical model that involves three major components in infection: resource assimilation of host, immune response of host and within-host dynamics of the pathogen. We propose that a feedback loop among those three components governs the progression and consequence of disease and infection and can yield a bistable outcome.

## MATERIAL AND METHODS

### A model of resource flow within the host

Here, we build a model based on the flow of within-host energetics/resources (thereafter resources) to understand infection dynamics. This view is motivated by several recent empirical studies [6,35–37]. For the host to maintain its energy and resource balance during infection, we assume that host’s intrinsic resources are drained in three ways: basic metabolism, immune response, and pathogen replication. Upon infection, host switches metabolism to distribute the energy and resources to combat pathogens [16] (Fig. 1).

**Fig. 1.**
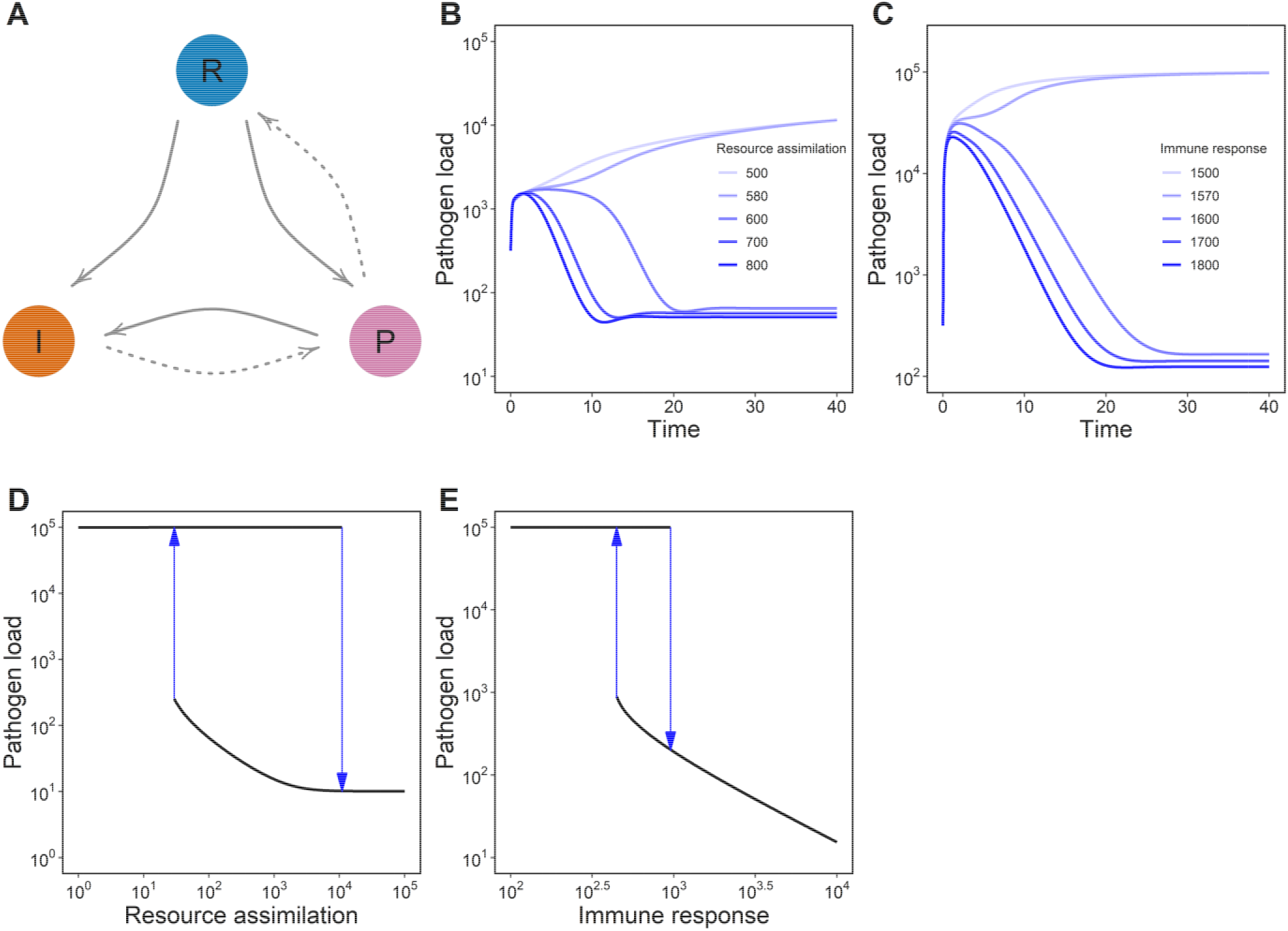
The bistability or hysteresis of host-pathogen interaction. (A) A schematic illustration of a minimal model which encapsulates within-host interactions of resources, immunity, and pathogens during the course of an infection. Solid arrows indicate positive interactions, while dashed arrows represent negative interactions. In such interaction with feedback loops, the pathogen population either propagates to the maximal carrying capacity of the host, or is reduced to a persistent low level dependent on resource assimilation, α (B), and immune response, β (C). Numerical simulations with varied initial values of P, R and I, show two distinct equilibria in terms of pathogen load across wide range of rate of resource assimilation (D) and Immune response (E). The four initial conditions are R = 10^0, I = 10^0, P = 10^4.99; R = 10^0, K = 10^0, P = 10^0; R = 10^6, K = 10^6, P = 10^4.99; R = 10^6, K = 10^6, P = 10^0.

In our model, hosts convert the ingested food, *C*, at an arbitrary rate, *α*, into instantaneously usable resources, *R*(see also Table S1 for all parameter definitions). The assimilation rate under infection is reduced by *ϕ*, which represents the virulence of pathogens: host assimilation under infection is *α*(1 − *ϕ*). We include this term of host assimilation under infection by assuming that bacterial endotoxins can block host metabolic functions, and hence host resource conversion and assimilation. Host resources, R, are used at rate *δ_R_* for maintaining metabolism, and at rate *λ* for immune defense. The host immune defense includes the production of immune effectors *I*, such as antimicrobial peptides and white blood cells, that use up resources. With the above assumptions, the rate of resource change can be written as

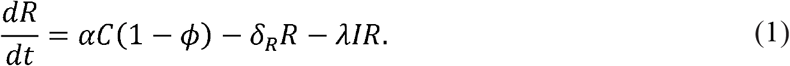

Once infected with pathogens, the host immune system responds to the infection by allocating resources to produce immune effectors *I*. The absolute quantity of immune factors scales to the pathogen density. We link the production rate of immune factors, *β*, to resource *R* and pathogen load *P*, by *βRPI*. At the same time, immune effectors are also removed by decay, for example they are metabolized, with rate *δ_i_*. The rate of immune effector change is described as:

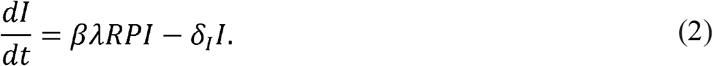

We first consider pathogen growth to be resource independent, but dependent on the carrying capacity inside the host. A more comprehensive interaction between host and pathogen will be discussed later. The rate of pathogen growth is described by the following relation,

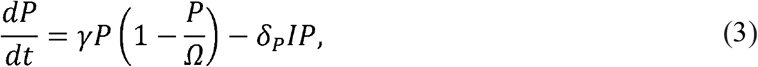

where *γ* is the arbitrary growth rate of pathogens, which is limited by the carrying capacity of the host *Ω* (if not by the availability of host resources). Pathogens are removed by the immune effectors with *δ_P_* and the natural death of pathogen can be neglected.

### The types of feedback interactions between pathogen and host

The assimilation and distribution of host resource are also constrained by the feedback of pathogens. Different pathogens may consume host resource to different extend. Some pathogens consume considerable amounts of host resources, such as parasitic wasps and bacteria in some insect hosts [15,17], or *Plasmodium* and gastrointestinal helminths in vertebrates [38]. Apart from eliciting immune responses, the propagation of pathogen and/or pathogen-related toxins can undermine the resource in-take or assimilation of the host, such as reducing foraging and slowing down metabolism, which subsequently will result in host death. Here we focus on host exploitation by pathogens consuming host resources or directly producing pathogenic factors to harm the host, we categorize the interactions between host and pathogen into three types: a pathogen consumes host resources and disrupt the host’s metabolic functions (A), a pathogen only consumes host resources (B), and a pathogen disrupts the host’s metabolic functions without consuming many resources (C).

In type A interactions the pathogen consumes host resources at rate *ϵ*. In addition, the pathogen disrupts host metabolism by reducing the assimilation of resource R, and the inhibition function *ϕ* is dependent on the pathogens; *ϕ* = *P*/(*P* + *Θ*_*P*→*R*_), where *P* is pathogen load and *Θ*_*P*→*R*_ is the half-saturation threshold of the pathogen that impairs the flow of resources inside the host. Experiments show that the immune response to pathogen infection dose does not increase proportionately with the pathogen load [35]. We model the immune response with respect to pathogen load by 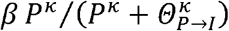. *κ* is a Hill coefficient, and *Θ*_*P*→*I*_ is the half saturation threshold of pathogen to elicit an immune response. Immune effectors, such as antimicrobial peptides, kill pathogens in a nonlinear fashion [45]. The death rate of pathogens has the form 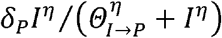. *η* is also a Hill coefficient, and *Θ*_*I*→*P*_ is the half saturation threshold of immune effectors to kill pathogens. Moreover, we assume that pathogen growth depends on the availability of resources as pathogens consume considerable resources. Modifying equation 1, equation 2 and equation 3 in the light of these assumptions, we obtain the full model of type A interaction with following dynamics:

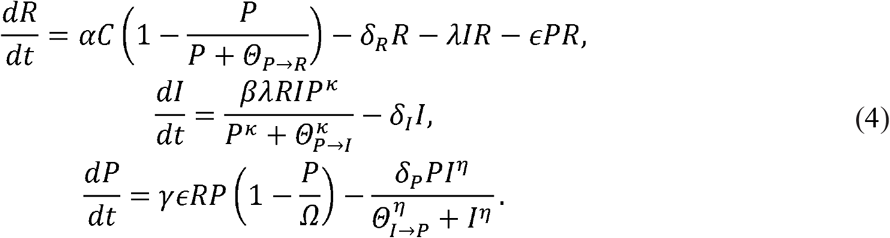

In Type B interaction, the host resource intake, *C*, and assimilation rate, *α*, are assumed independent of pathogen load as the pathogen causes little or no harm to the host metabolic functions. Pathogens consume resources from the host, and pathogen growth is thus resource dependent. Immune response and pathogens elimination also follow a Hill Function. For those pathogens that fall into this category, the host pathogen interaction can be captured by (also see Fig. 1 and Fig. S3):

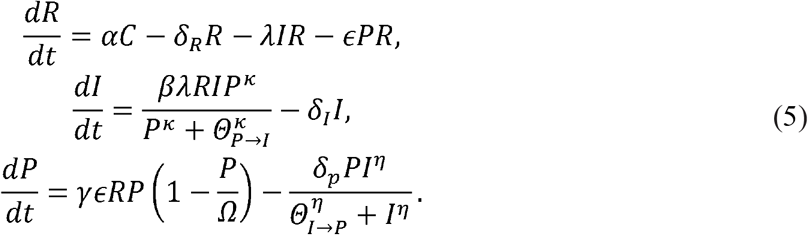

In Type C interaction, the pathogen sabotages host metabolic functions, but consumes little resources from the host. The growth of pathogen is therefore independent of host resources. With similar assumptions as above regarding immune responses and pathogen elimination, we use the following equations to capture the interaction:

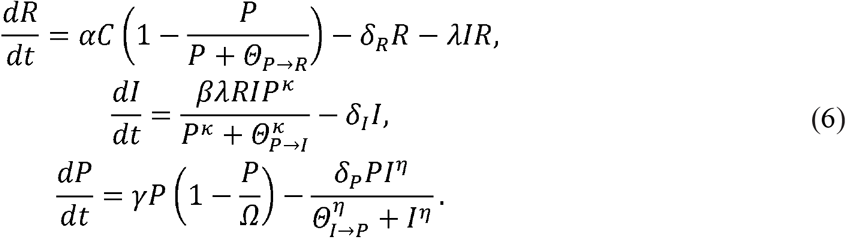

Consequently, if the host rapidly eliminates the pathogen or keeps the pathogen under threshold level and is symptom free, it will eventually survive, as long as basal metabolism does not exceed consumption and assimilation efficiency. Occasionally, the host could also maintain very high pathogen load while showing no or rather mild symptoms of infection which usually indicates high “tolerance” of the host. If pathogens are less resource costly, keeping large numbers of pathogens inside the body requires the host having an efficient turn-over of food into within host resources to clear toxic substance and repair host damage generated by pathogens. Thus, it is crucial for a host to balance the cost of maintaining high metabolic activity and reducing the damage of pathogen.

### Model analysis and simulation

Using the models described above, we analyzed how the different parameter settings determine the final infection outcome within a host. We solve the equations numerically with sets of parameters and initial conditions (table S1). To examine the bistable infection outcomes of host individuals, we simulated dynamics of infection in individuals with four varied initial conditions (R = 10^0, I = 10^0, P = 10^4.99; R = 10^0, K = 10^0, P = 10^0; R = 10^6, K = 10^6, P = 10^4.99; R = 10^6, K = 10^6, P = 10^0).

To explore the criticality and sensitivity of the interaction, we numerically solved the system until the steady states are reached. The steady states are identified when the difference of solutions between two time points are less than 10^-5. We then record the pathogen load at the end of each simulation and analyzed the bistability across simulations. To broadly explore how the initial settings of resource level, immunity and pathogen load affect the infection outcomes, we set an orthogonal combination of three initial conditions ranging from 10^0 to 10^5. We then identified two infection trajectories by monitoring the pathogen load. If the pathogen load reaches the host’s carrying capacity, the trajectory is classed as “death”, otherwise as “survival”.

To investigate how intrinsic noise affect the dynamics of infection, we ran simulations by adding a noise term to each of the states in the deterministic model. For each simulation we drew one random number from a normal distribution with mean 0 and variance of σ^2^. We ran 500 simulations for each noise level, and explored noise levels between values of 10^-5 to 10^-2. We repeated these sets of simulations for each of the three initial parameter settings, resource assimilation, immune response, and pathogen load.

In order to simulate the bimodal distribution of pathogen load in experiments, we ran simulations by drawing the parameter value of resource assimilation from a uniform distribution *α*~U[1500, 1700], and the parameter value of immune response from a logarithmic uniform distribution *λ*~10^U[2, 5]. Such draws were conducted for 5000 samples. We then recorded the pathogen load at different time points in each replicate. All the calculations and simulations are implemented in R 3.6.0.

## RESULTS

### Bistable infection outcomes

We modeled the in-host infection dynamic of simple interactions among three components: the flow of within-host resources, immune response, and pathogen load (Fig. 1a). Our analysis showed that the interactions of these three components collectively determines the outcome of the infection. In particular, disease progression falls into two different trajectories: (a) pathogen load increases to the maximum and (b) a decline of pathogen load to some controlled level that resembles a chronic infection (Fig. 1b, Fig. 1c). This pattern reveals a typical pattern of bistability. Resource supply, immune activation, and pathogen load, follow two distinct trajectories which determine death or survival of the host (Fig. S1, Fig. S2).

Our analysis also showed that host resource assimilation and the strength of the immune response are two important contributors for the bistable pattern of an infection (Fig. 1d, Fig. 1e). If the rate of resource assimilation and immune response is lower than the threshold level, the pathogen population will grow to high titres that can lead to the death of the host. If the rate of resource assimilation and immune response is at intermediate levels, the pathogen either propagates or is controlled. If the rate of resource assimilation and immune response is above threshold levels, the pathogen will always be controlled (Fig.1d, Fig. 1e).

Different pathogens manipulate hosts in different ways by controlling the regulatory relations among host resource assimilation, host immune response, and pathogen virulence. Our analysis further revealed that the ways how different pathogens manipulate hosts does not qualitatively change the bistable outcome of the host pathogen interaction (Fig. S3).

### Initial levels of resource, immunity and pathogen load before infection determine the infection trajectory

Immune challenge and environmental stress can alter reservoirs of host resources and levels of host immunity prior to infection. One would also expect that immune challenged hosts having a higher level of basic immunity are likely to survive another infection. We investigated whether changes or variations in initial host immunity and resource reservoir conditions together with pathogen load would impact the outcome of infection. By fixing the rates of resource assimilation and immune reaction, we find that the dynamics of infection are sensitive to the initial conditions of level of immunity, the availability of resources, and pathogen load before infection (Fig. 2a, Fig 2b). Sampling across a wide range of the three initial parameters—immune response, pathogen load and host resources — in our model, we found that increased initial immunity increased host survival, higher initial pathogen load increased host mortality, however, the initial resource reservoir did not affect survival probabilities(Fig 2c). We assessed combined effects of initial parameter settings and thereby revealed that small differences in quantity of the combinations can result in distinct infection trajectories (Fig. S4) in certain ranges.

**Fig. 2.**
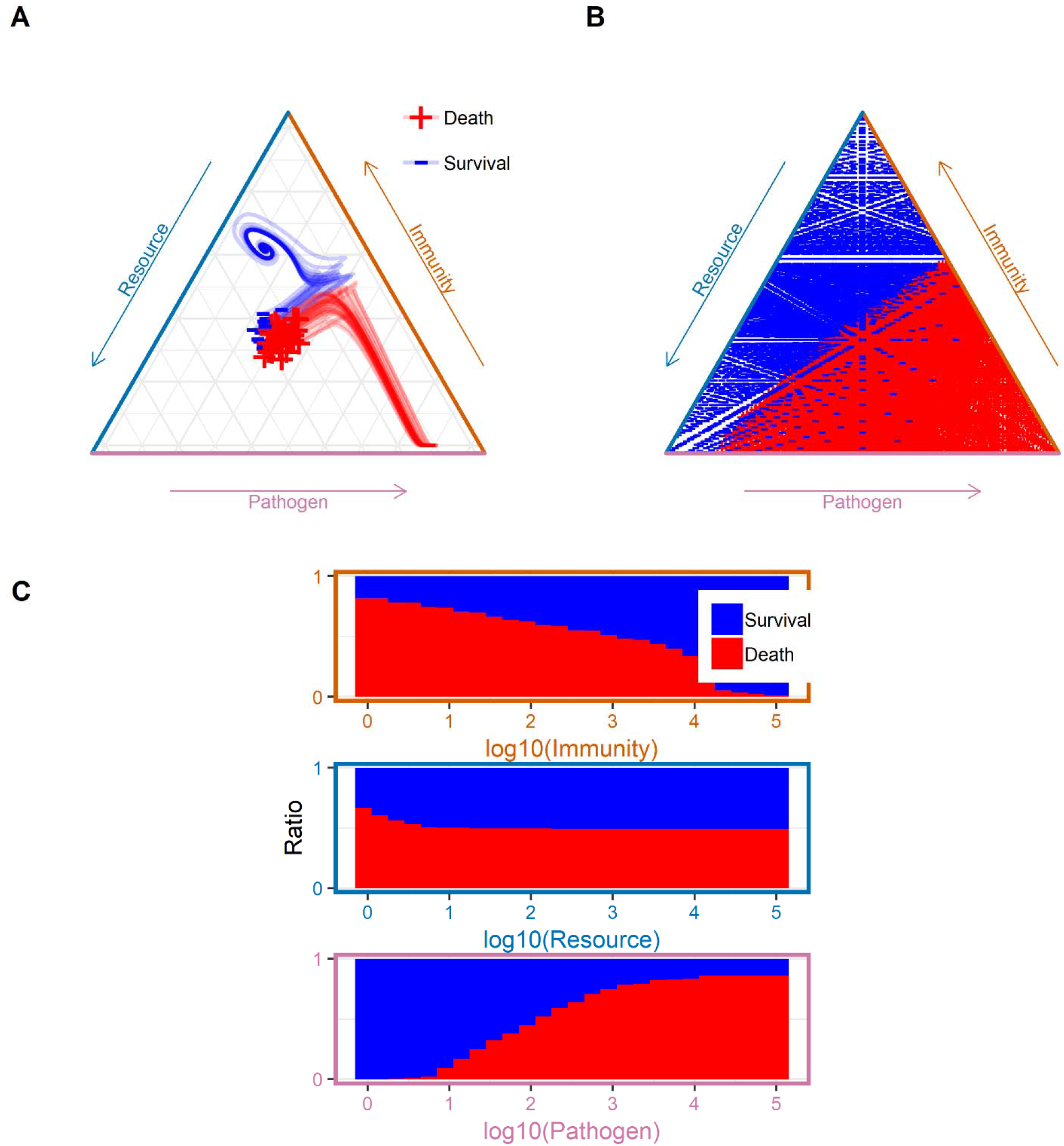
Small variations in initial conditions determine distinct infection outcomes. (A, B) With identical rate parameters, small differences in initial quantity of resource, immunity, and pathogen load will lead to different disease trajectories during infection. (C) The pooled distribution of trajectories of death and survival against initial parameter settings of immunity, resource, and pathogen. This suggests that a wide range of initial level of these “state” will cause bimodal distribution of death and survival as the consequence of an infection. The range of initial conditions for resource, immunity and pathogen are all from 10^0 to 10^5.

### Intrinsic noise fosters bistability

We next examined whether intrinsic noise leads to bistable infection outcomes. First, we fixed the deterministic system at a state that pathogen will not be cleared. Then, we introduced a noise term either into resource assimilation, immune reaction, or pathogen growth. We found that when the noise level is low, the system will maintain the previously set state, in which the infection is acute and the pathogen will not be cleared. As the noise level increase, the system either falls into the controlled state, in which pathogen will be kept at low abundance and the host suffers chronic infection, or into the uncontrolled state, in which the pathogen will grow and the host suffers acute infection. In addition, our results show that noise in resource assimilation, immune response, and pathogen growth all contribute to bistability in the infection outcomes at the critical level (Fig. 3).

**Fig. 3.**
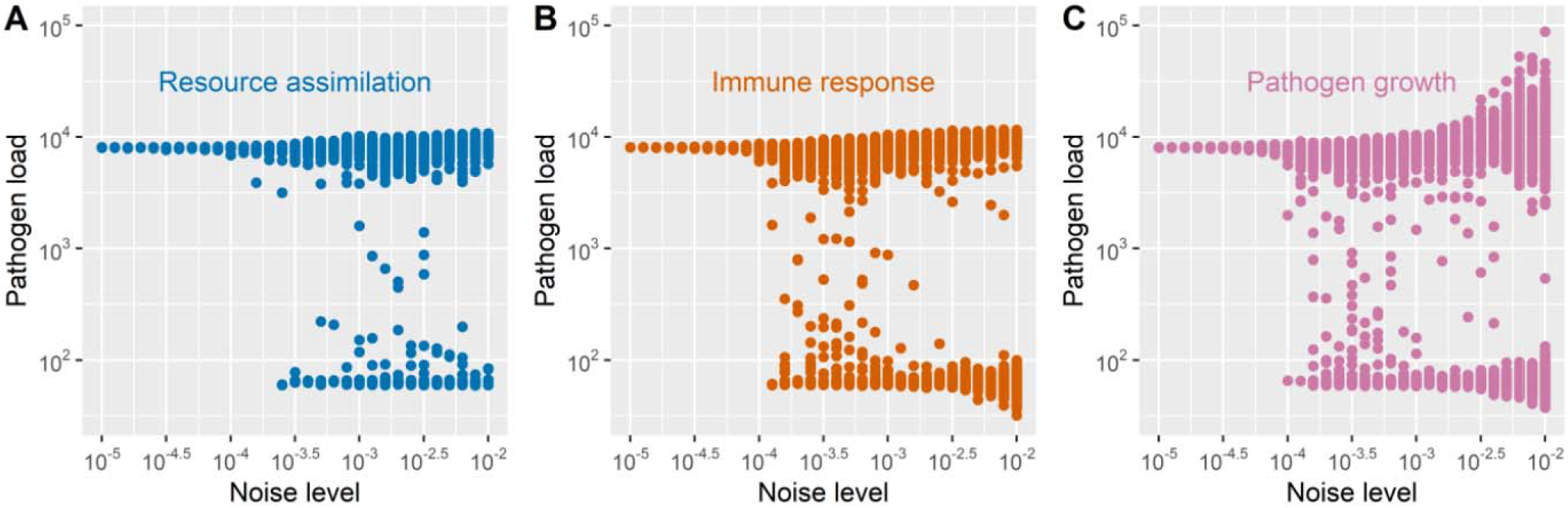
Intrinsic stochasticity or noise results in bistable infection outcomes. Perturbation of the dynamics of infections changes the disease trajectories which resulted in bimodal distributions of pathogens. When the noise level is low, all cases of infection have only one equilibrium in which infection is acute and the pathogen load is high. As the noise increase, infection in some cases will be controlled, in which the pathogen load is maintained lowly. Moreover, intrinsic noise in resource (A), immunity (B) and pathogen growth (C) will all cause the same patterns of bimodal distribution.

### Bistability leads to variations in pooled time-series data

In Fig. 4 we simulated the progression of infection and *in-silico* sampled the bacterial load at different time points in host individuals. In the pooled data, bacterial load diversified after initial infection, reached considerably variable bacteria load at some intermediate time points during progression, before finally converging to a bimodal distribution. Variations in bacterial load create differences with several orders of magnitude. We then explored two different parameters settings: a) fixed initial conditions with varied rates of resource assimilation and immune response, and b) varied initial condition with fixed rates of resource assimilation and immune response. In both cases, the pathogen load in hosts sampled at different time points first followed the same pattern, a unimodal distribution, followed by a bifurcation, i.e. splitting into a bimodal distribution (Fig 4).

**Fig. 4.**
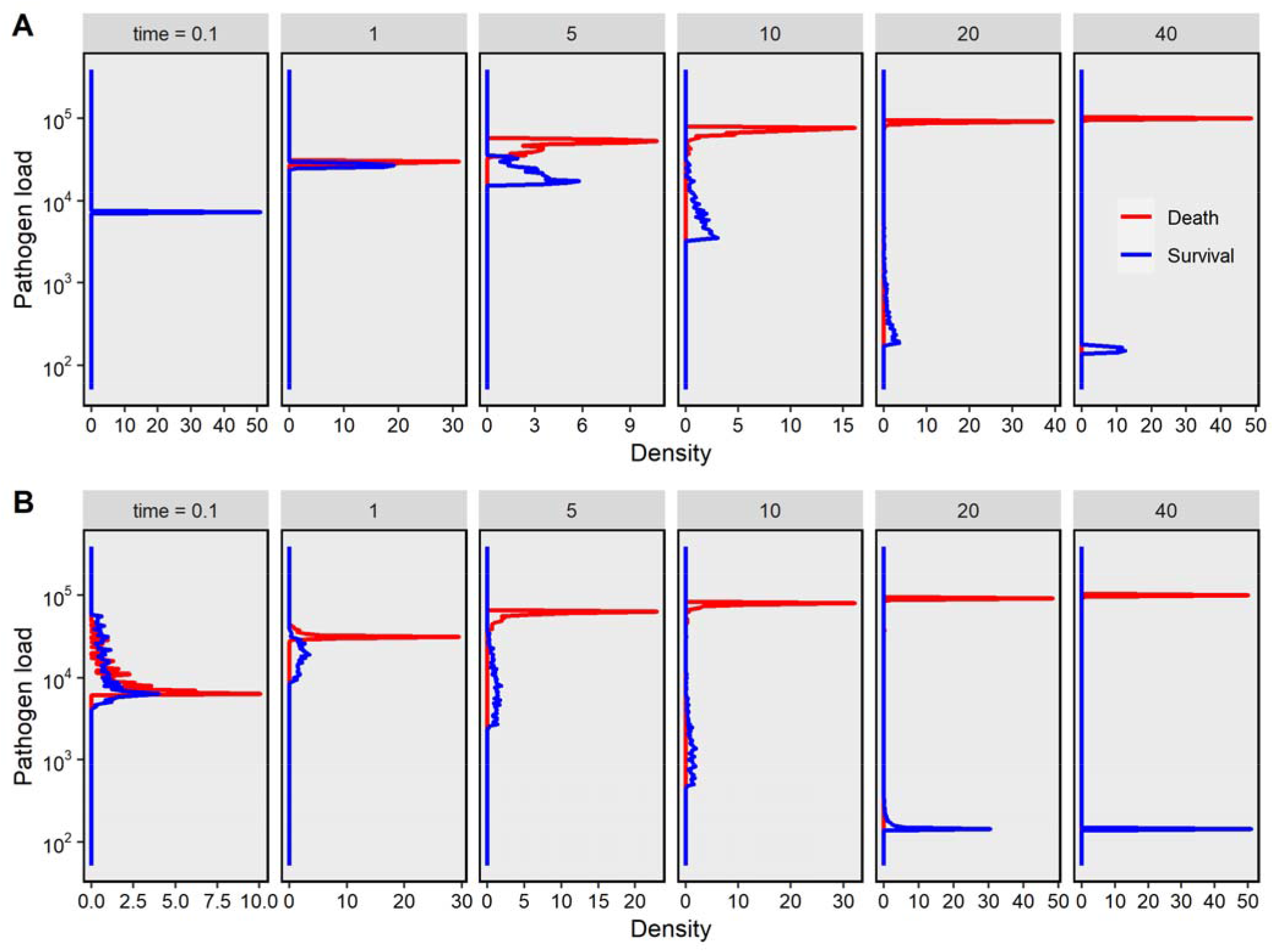
Bistability leads to variations in pooled time-series data. We numerically solve the model with varied immune response rates (A) and initial quantity of pathogens (B), in which the distributions of both parameters are following the uniform distribution α~U[1500, 1700] and logarithmic uniform distribution β~10^U[2, 5], respectively. Data sampled from different time points shows large variations with several orders of magnitude.

## DISCUSSION

In this study, we modeled a feedback in host resource assimilation, host immune response and pathogen performance. These parameters interacted and generated bistable infections that lead to either survival or death of the host. In particular, the bistability of our model depends on the rates of immune response and metabolism as it is highly sensitive to the rate parameters, α and β. This may be the result of collective effects of the underlying molecular mechanisms, for example, spontaneous gene up-regulation during infection could shift hosts to the survival trajectory [16]. In addition, immune responses, such as the expression of antimicrobial peptides, tend to show correlated spatial and temporal patterns during infection [16]. Moreover, a recent study investigating a densely sampled time-series of whole genome transcriptomics in *Drosophila* found that genes in the same “functional clusters” displayed correlated expression patterns during the course of infection [40]. Also, pathogens can manipulate the host response by reducing the resource assimilation of the host, a reduction in food intake and compromised metabolism [17,35,41]. Reduced resource assimilation subsequently lowers host condition and decreases the ability to maintain homeostasis and to clear infections. This is partially due to toxic factors released by pathogens. Pathogens producing no toxic factors, but directly competing for resources with the host, may also compromise host condition and can result in the same bistable outcome (Fig S3). In line with previous theoretical studies that highlight resource availability for fighting and tolerating pathogens [17,35,41], our simple model captures the feedbacks among immunity, pathogens, and host resources and explains the survival and death of host in an infection. We highlight that the feedbacks among host resource, pathogen load, and the ability to maintain immunity are potentially key mechanisms to determine the infection outcome.

Our model found that bistability of host-pathogen interactions is very sensitive to initial conditions. For example the baseline of immunity and resources, are critical for determining the bistable infection outcome. Variations of baseline immunity, resource, and pathogen load can be the result of differential food supply or prior pathogen exposure. Changes in baseline immunity are commonly observed in studies on “memory effects” of invertebrate immune systems [42,43]. Although our model, together with published experimental work, finds that higher levels of baseline immunity lead to better infection outcomes, there is still large uncertainty and variation at the individual level. Slight differences in the combination of initial conditions (immunity, resource, and pathogen load) will lead to distinct trajectories of life and death (Fig. 2A, 2B). It is known that most immune challenges up-regulate cellular and humoral immunity, but it probably changes levels of other physiological aspects such as carbohydrate and amino acids resources, before the whole system reverts to homeostasis [44]. To observe the divergent infection outcomes, we propose that future experiments could be designed to monitor a variety of related physiological states with reasonable time-resolution, which would also require large sample sizes [31,40,44].

Our model also finds that pathogen load sampled from different host individuals at different time points varies substantially, a finding commonly observed in experiments [15,27,31,35,45–49]. We reason that this variation could result from the bistable characteristics. In large animal models, for example following *Plasmodium* infection in individual mice revealed distinct damped oscillating pattern of pathogen load within the host [27,50]. However, for smaller model hosts such as *Drosophila*, continuously monitoring pathogens dynamics in the same individuals is largely constrained. To circumvent these constraints, researchers sampled pooled times series data from different individuals at the same time point [31,45]. This causes large variations in pathogen distribution even in precisely controlled experimental conditions. Sampling pathogen loads with finer time-resolution revealed bifurcating pathogen loads and their consequence of bistable outcome. Our model seeks to explain this phenomenon. We suggest that the nonlinear characteristics of the host-pathogens interaction might be sufficient to generate these bistable outcomes. Individual hosts could progress along different infection trajectories, for which pathogen load varies at single time points. We further examined the stochastic nature within the host and found that small internal perturbations of immune responses and resource metabolism could drive the infection from the one steady state to another.

We also investigated how different type of pathogens influence the patterns of infection. Pathogens, such as viruses, may cause severe harm to hosts, but consume little resources of hosts [51]; alternatively, they may cause little direct harm to the host, but consume large amount of resources of hosts, for example in the case of helminth [52]. We separately implemented these distinct pathogens in our model. Our results showed that the bistable nature of the infection dynamics does not qualitatively change for different pathogens in our models.

In our model, solutions of states co-determined the trajectories of infection outcomes and recovered the diagram of disease trajectories. Relating the change of pathogen load to the host’s health is crucial to understand the disease progression and provides insight into understanding concepts such as tolerance vs. resistance. Researchers have shown a correlation between host health and pathogen load in time-series data [25,30]. These correlations shape different trajectories on the “phase plane” that differentiate survival and death of hosts. This is consistent with the idea that infection outcomes in various circumstance are bistable. Moreover, correlating pathogen load with many other physiological and immunological states essentially generate similar patterns. In short, many infection-related physiological states together shape the nonlinearity of the in-host infection dynamics. Our model can provide a mechanistic explanation for the distinct infection trajectories of the death-survival diagram [25,30](see also Fig S4).

We revealed the nonlinear interaction between host and pathogen, in which disease progression followed two distinct trajectories which lead to death and survival of hosts. The catastrophic shift in some critical parameters determines the death and survival of the host. Thus, it is crucial to identify these critical conditions in which a catastrophic shift will happen. This includes tracking pathogen load, host immunity, and nutrition level in real time. Treating infectious disease is to probe the non-linearly dynamical system, and trying to delay the catastrophic shift by manipulating particular parameters. Besides directly killing pathogens, these strategies also include using drugs to neutralize the toxic substance secreted by the pathogens, which could substantially reduce the damage to the host and increase host tolerance. Moreover, metabolic dysfunction, caused by infection, will drastically reduce the host’s ability to maintain homeostasis during infection. Extra supplement of energy may eventually help overcoming the infection. These properties that emerge from a non-linear system should be taken into consideration when developing a more comprehensive approach to treat infections.

## Note

At the final stage of writing this manuscript, a related work was brought to our attention: Stephen P. Ellner, Nicolas Buchon, Tobias Dörr, Brian P. Lazzaro. Host-pathogen Immune Feedbacks Can Explain Widely Divergent Outcomes from Similar Infections. bioRxiv. doi: https://doi.org/10.1101/2021.01.08.425954

## Acknowledgments

We thank Dr. Alexandro Rodríguez-Rojas, Dr. Sophie Armitage and Dr. Désirée Bäder for inspiring discussion at the early stage of this work.

---

**Fig. S1.**
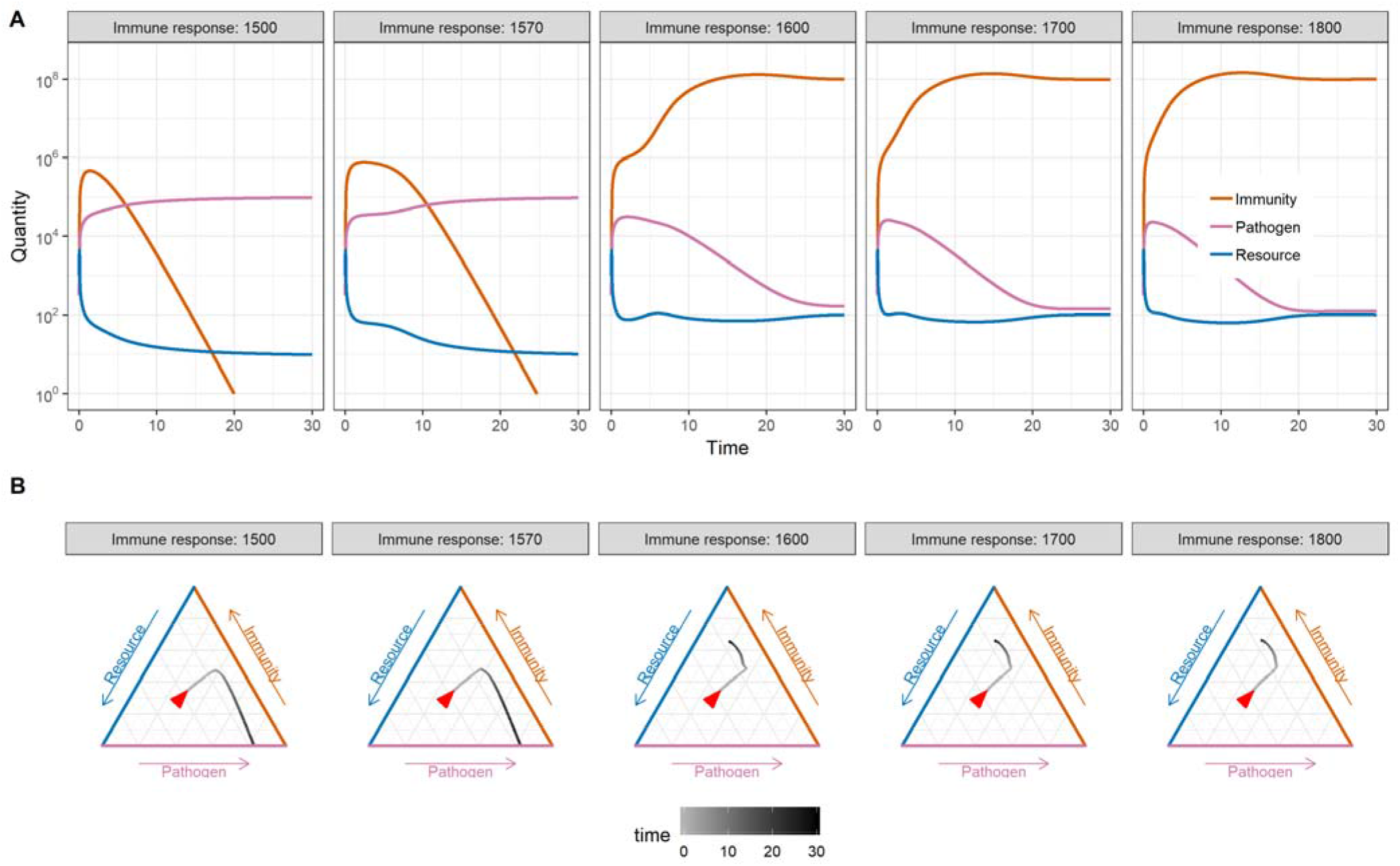
The dynamics of immunity, pathogen load, and resource levels for different levels of immune response (A). These patterns fall into two different trajectories (B).

**Fig. S2.**
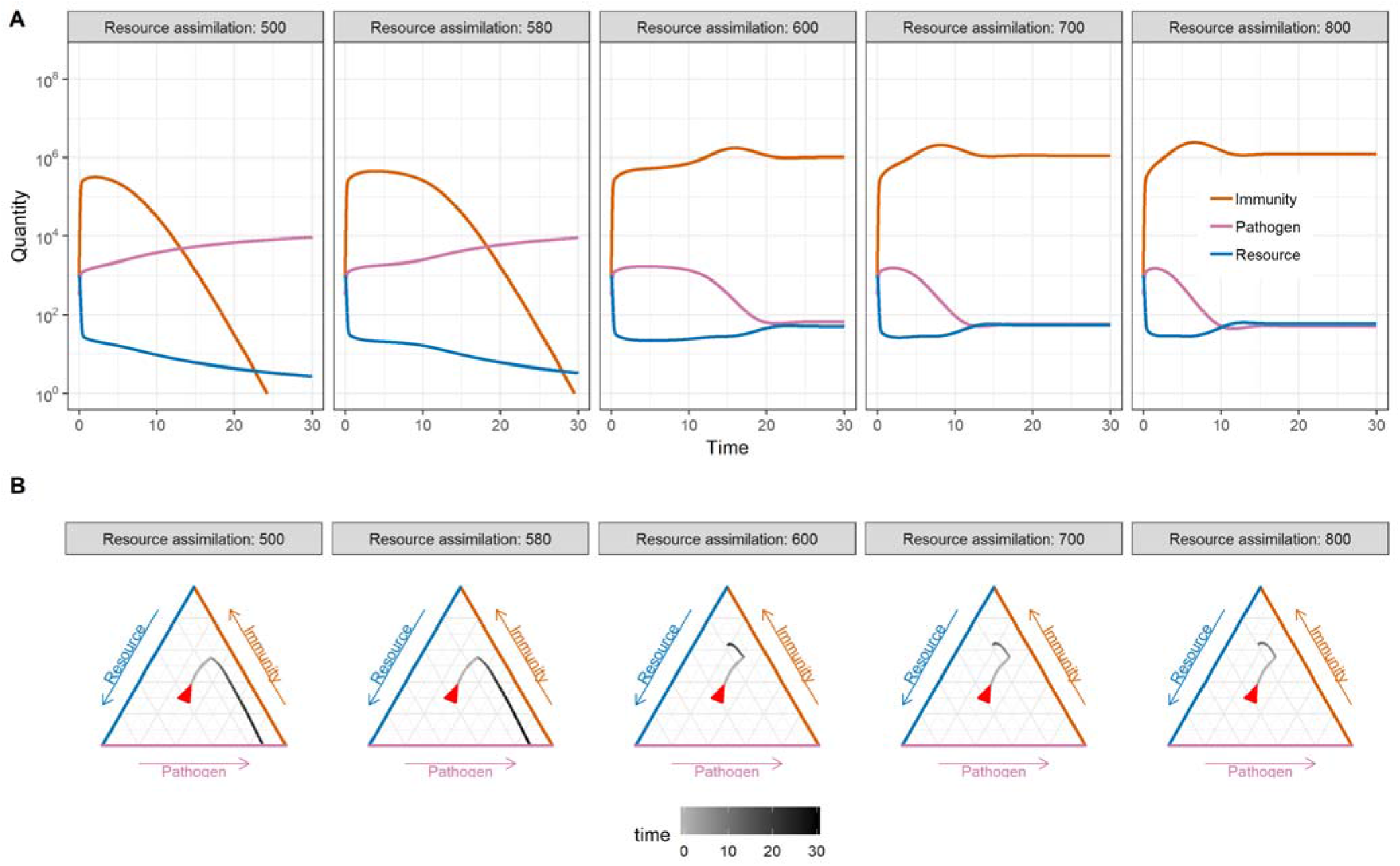
The dynamics of immunity, pathogen load and resources levels showed different patterns under different levels of resource assimilation (A). These patterns fall into two different trajectories (B).

**Fig. S3.**
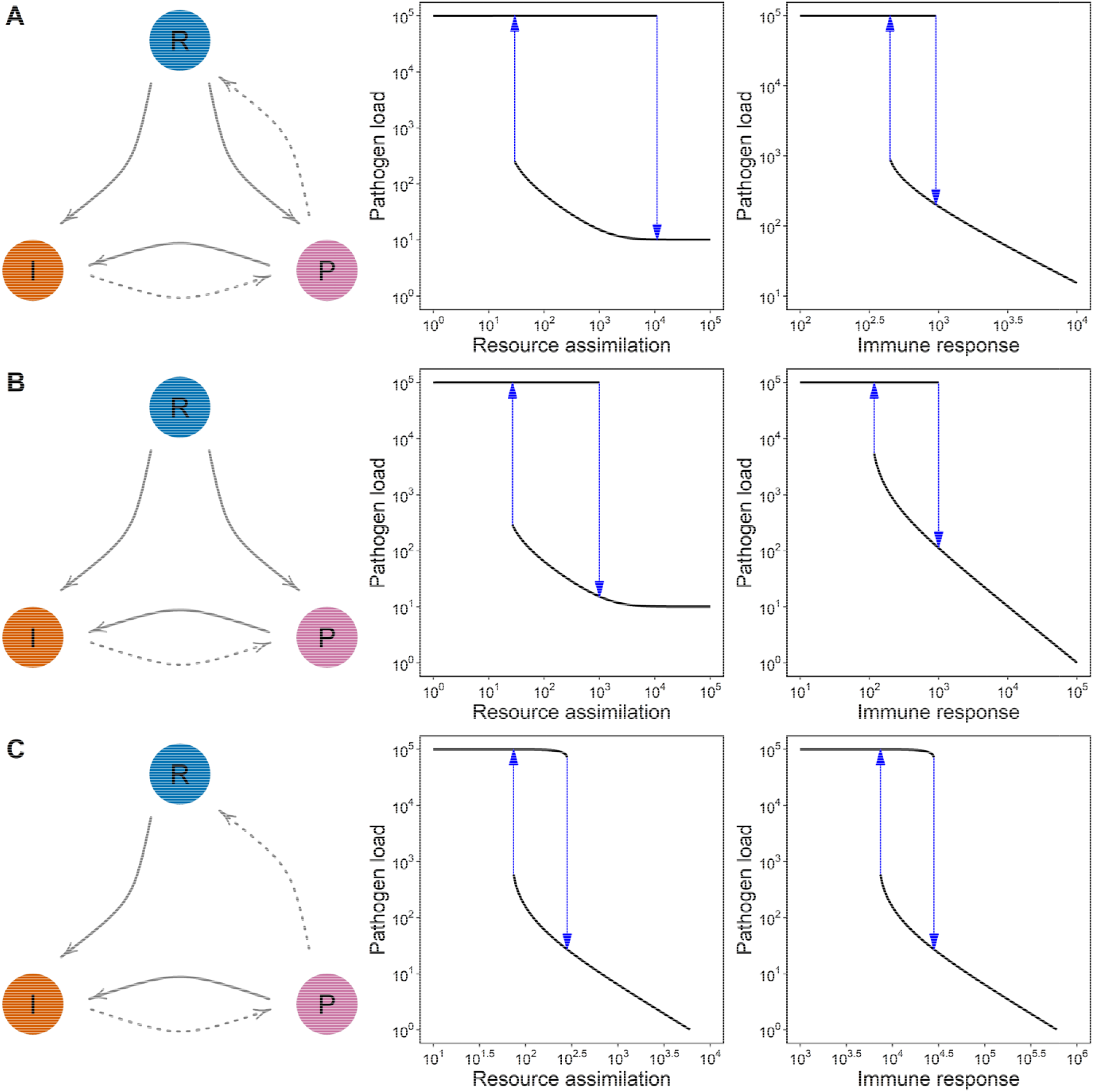
Infections with different patterns of in-host interaction share the bistability with respect to resource assimilation and immune response.

**Fig. S4.**
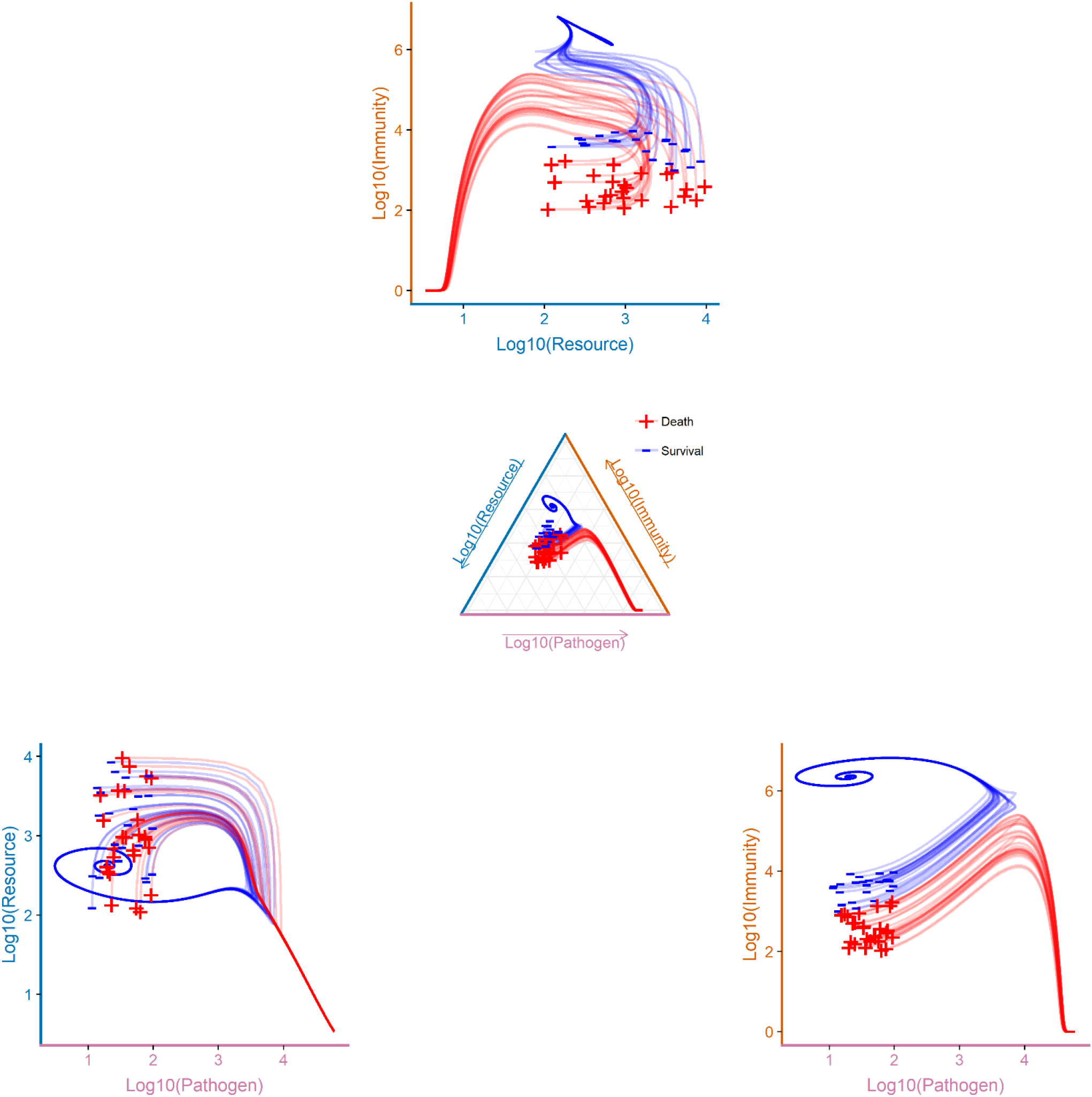
The infection trajectories are resembled by two-by-two correlation of resource level, immunity and pathogen load, in which there are always two trajectories representing the consequences of the infection. Red means host killed by pathogens; blue means host overcoming pathogens either killing them all or survive with persistence of low pathogen load.

**Table S1.** Variables and parameters used in the model. All the default values, if not specified elsewhere, are listed below.

| Variables/Parameters | Default values | Description |
| --- | --- | --- |
| $R$ | $10^3$ | Total directly usable resource in host, such as carbohydrates and amino acids. |
| $P$ | $10^{2.5}$ | Pathogen load. |
| $I$ | $10^3$ | Overall immunity level, which includes blood cells related to immune function, immune peptides phenoloxidases and other related antipathogen enzymes. |
| $\Omega$ | $10^5$ | Physiological and physical carrying capacity of pathogens in host. |
| $\Theta_{P \rightarrow R}$ | $10^4$ | Threshold values of pathogens for affecting resource assimilation. |
| $\Theta_{P \rightarrow I}$ | $10^2$ | Threshold values of pathogens for initiating immune response. |
| $\Theta_I$ | $10^6$ | Threshold values of immunity for killing pathogens. |
| $C$ | 1 | Total in-take food. |
| $\alpha$ | $10^4$ | Rate of resource assimilation. |
| $\beta$ | 3000 | Rate of immune response, for example, the rate of generating immune substance. |
| $\gamma$ | 1 | Growth rate of pathogen. |
| $\delta_R$ | 1 | Rate of resources consumption by host. |
| $\delta_I$ | 1 | Clearing rate of immune substance. |
| $\delta_P$ | 1 | Clearing rate of pathogen. |
| $\varepsilon$ | $10^{-2}$ | Rate of resource consumption of pathogen. |
| $\lambda$ | $10^{-5}$ | Rate of resource consumption of immunity. |
| $\kappa$ | 1 | Hill coefficient for pathogens. |
| $\eta$ | 2 | Hill coefficient for immunity. |

## References

1. Haine, E. R., Moret, Y., Siva-Jothy, M. T., Rolff, J. 2008 Antimicrobial defense and persistent infection in insects. Science 322, 1257–1259. (10.1126/science.1165265)

2. Jackson, J. A., Hall, A. J., Friberg, I. M., Ralli, C., Lowe, A., Zawadzka, M., Turner, A. K., Stewart, A., Birtles, R. J., Paterson, S., Bradley, J. E., Begon, M. 2014 An immunological marker of tolerance to infection in wild rodents. Plos Biol. 12, e1001901. (10.1371/journal.pbio.1001901)

3. Lynn, B. M. I., Scheuerlein, A., Wikelski, M. 2003 Immune activity elevates energy expenditure of house sparrows: A link between direct and indirect costs? Proc. Roy. Soc. B. 270, 153–158. (10.1098/rspb.2002.2185)

4. Rolff, J., Schmid-Hempel, P. 2016 Perspectives on the evolutionary ecology of arthropod antimicrobial peptides. Phil. Trans. Roy. Soc. B 371, 20150297. (10.1098/rstb.2015.0297)

5. Kutzer, M. A. M., Armitage, S. A. O. 2016 The effect of diet and time after bacterial infection on fecundity, resistance, and tolerance inDrosophila melanogaster. Ecol. Evol. 6, 4229–4242. (10.1002/ece3.2185)

6. Cumnock, K., Gupta, A. S., Lissner, M., Chevee, V., Davis, N. M., Schneider, D. S. 2018 Host energy source is important for disease tolerance to malaria. Curr. Biol. 28, 1635–1642. (10.1016/j.cub.2018.04.009)

7. Galenza, A., Hutchinson, J., Campbell, S. D., Hazes, B., Foley, E. 2016 Glucose modulates Drosophila longevity and immunity independent of the microbiota. Biol. Open 5, 165–173. (10.1242/bio.015016)

8. Becker, D. J., Streicker, D. G., Altizer, S. 2015 Linking anthropogenic resources to wildlife-pathogen dynamics: A review and meta analysis. Ecol. Lett. 18, 483–495. (10.1111/ele.12428)

9. Ponton, F., Wilson, K., Holmes, A. J., Cotter, S. C., Raubenheimer, D., Simpson, S. J. 2013 Integrating nutrition and immunology: A new frontier. J. Insect Phys. 59, 130–137. (10.1016/j.jinsphys.2012.10.011)

10. Cotter, S. C., Littlefair, J. E., Grantham, P. J., Kilner, R. M. 2013 A direct physiological trade-off between personal and social immunity. J. Anim. Ecol. 82, 846–853. (10.1111/1365-2656.12047)

11. Budischak, S. A., Hansen, C. B., Caudron, Q., Garnier, R., Kartzinel, T. R., Pelczer, I., Cressler, C. E., van Leeuwen, A., Graham, A. L. 2017 Feeding immunity: Physiological and behavioral responses to infection and resource limitation. Front. Immunol. 8, 1914. (10.3389/fimmu.2017.01914)

12. Unckless, R. L., Rottschaefer, S. M., Lazzaro, B. P. 2015 The complex contributions of genetics and nutrition to immunity in drosophila melanogaster. Plos Genet. 11, e1005030. (10.1371/journal.pgen.1005030)

13. Wang, A., Huen, S. C., Luan, H. H., Baker, K., Rinder, H., Booth, C. J., Medzhitov, R. 2018 Glucose metabolism mediates disease tolerance in cerebral malaria. Proc. Natl. Acad. Sci. U.S.A.. 115, 11042–11047. (10.1073/pnas.1806376115)

14. Povey, S., Cotter, S. C., Simpson, S. J., Wilson, K. 2014 Dynamics of macronutrient self medication and illness induced anorexia in virally infected insects. J. Anim. Ecol. 83, 245–255. (10.1111/1365-2656.12127)

15. Ayres, J. S., Schneider, D. S. 2009 The role of anorexia in resistance and tolerance to infections in drosophila. Plos Biol. 7, e1000150. (10.1371/journal.pbio.1000150)

16. Sanchez, K. K., Chen, G. Y., Schieber, A. M. P., Redford, S. E., Shokhirev, M. N., Leblanc, M., Lee, Y. M., Ayres, J. S. 2018 Cooperative metabolic adaptations in the host can favor asymptomatic infection and select for attenuated virulence in an enteric pathogen. Cell 175, 146–158. (10.1016/j.cell.2018.07.016)

17. Bajgar, A., Kucerova, K., Jonatova, L., Tomcala, A., Schneedorferova, I., Okrouhlik, J., Dolezal, T. 2015 Extracellular Adenosine Mediates a Systemic Metabolic Switch during Immune Response. Plos Biol. 13, e1002135. (10.1371/journal.pbio.1002135)

18. Lazzaro, B. P., Sackton, T. B., Clark, A. G. 2006 Genetic variation in drosophila melanogaster resistance to infection: A comparison across bacteria. Genetics 174, 1539–1554. (10.1534/genetics.105.054593)

19. Unckless, R. L., Lazzaro, B. P. 2016 The potential for adaptive maintenance of diversity in insect antimicrobial peptides. Phil. Trans. Roy. Soc B 371, 20150291. (10.1098/rstb.2015.0291)

20. Hotson, A. G., Schneider, D. S. 2015 Drosophila melanogaster Natural Variation Affects Growth Dynamics of InfectingListeria monocytogenes. G3 Genes/Genomes/Genetics 5, 2593–2600. (10.1534/g3.115.022558)

21. Vogel, H., Müller, A., Heckel, D. G., Gutzeit, H., Vilcinskas, A. 2018 Nutritional immunology: Diversification and diet-dependent expression of antimicrobial peptides in the black soldier fly *Hermetia illucens*. Dev. Comp. Immunol. 78, 141–148. (10.1016/j.dci.2017.09.008)

22. Little, T. J., Shuker, D. M., Colegrave, N., Day, T., Graham, A. L. 2010 The coevolution of virulence: Tolerance in perspective. Plos Pathog 6, e1001006. (10.1371/journal.ppat.1001006)

23. Ayres, J. S., Schneider, D. S. 2012 Tolerance of infections. Annu. Rev. Immunol. 30, 271–294. (10.1146/annurev-immunol-020711-075030)

24. Råberg, L. 2014 How to live with the enemy: Understanding tolerance to parasites. Plos Biol. 12, e1001989. (10.1371/journal.pbio.1001989)

25. Torres, B. Y., Oliveira, J. H. M., Thomas Tate, A., Rath, P., Cumnock, K., Schneider, D. S. 2016 Tracking resilience to infections by mapping disease space. Plos Biol. 14, e1002436. (10.1371/journal.pbio.1002436)

26. Graham, A. L., Tate, A. T. 2017 Are we immune by chance? Elife 6. (10.7554/eLife.32783)

27. Mideo, N., Savill, N. J., Chadwick, W., Schneider, P., Read, A. F., Day, T., Reece, S. E. 2011 Causes of variation in malaria infection dynamics: Insights from theory and data. Am. Nat. 178, E174–E188. (10.1086/662670)

28. Louie, A., Song, K. H., Hotson, A., Thomas Tate, A., Schneider, D. S. 2016 How many parameters does it take to describe disease tolerance? Plos Biol. 14, e1002435. (10.1371/journal.pbio.1002435)

29. Wale, N., Sim, D. G., Jones, M. J., Salathe, R., Day, T., Read, A. F 2017 Resource limitation prevents the emergence of drug resistance by intensifying within-host competition. Proc. Natl. Acad. Sci. U.S.A.. 114, 13774–13779. (10.1073/pnas.1715874115)

30. Lough, G., Kyriazakis, I., Bergmann, S., Lengeling, A., Doeschl-Wilson, A. B. 2015 Health trajectories reveal the dynamic contributions of host genetic resistance and tolerance to infection outcome. Proc. Roy. Soc. B. 282, 20152151. (10.1098/rspb.2015.2151)

31. Duneau, D., Ferdy, J., Revah, J., Kondolf, H., Ortiz, G. A., Lazzaro, B. P., Buchon, N. 2017 Stochastic variation in the initial phase of bacterial infection predicts the probability of survival in D. Melanogaster. Elife 6. (10.7554/eLife.28298)

32. Cressler, C. E., Graham, A. L., Day, T. 2015 Evolution of hosts paying manifold costs of defence. Proc. Roy. Soc. B. 282, 20150065. (10.1098/rspb.2015.0065)

33. Frank, S. A. 2002 Immune response to parasitic attack: Evolution of a pulsed character. J. Theor. Biol. 219, 281–290. (10.1006/jtbi.2002.3122)

34. Tate, A. T., Graham, A. L. 2015 Dynamic patterns of parasitism and immunity across host development influence optimal strategies of resource allocation. Am. Nat. 186, 495–512. (10.1086/682705)

35. van Leeuwen, A., Budischak, S. A., Graham, A. L., Cressler, C. E. 2019 Parasite resource manipulation drives bimodal variation in infection duration. Proc. Roy. Soc. B. 286, 20190456. (10.1098/rspb.2019.0456)

36. Cressler, C. E., Nelson, W. A., Day, T., McCauley, E. 2014 Disentangling the interaction among host resources, the immune system and pathogens. Ecol. Lett. 17, 284–293. (10.1111/ele.12229)

37. Canale, C. I., Henry, P. 2011 Energetic costs of the immune response and torpor use in a primate. Funct. Ecol. 25, 557–565. (10.1111/j.1365-2435.2010.01815.x)

38. Budischak, S. A., Wiria, A. E., Hamid, F., Wammes, L. J., Kaisar, M. M. M., van Lieshout, L., Sartono, E., Supali, T., Yazdanbakhsh, M., Graham, A. L. 2018 Competing for blood: The ecology of parasite resource competition in human malaria-helminth co-infections. Ecol. Lett. 21, 536–545. (10.1111/ele.12919)

39. Johnston, P. R., Makarova, O., Rolff, J. 2014 Inducible defenses stay up late: Temporal patterns of immune gene expression inTenebrio molitor. G3 Genes/Genomes/Genetics 4, 947–955. (10.1534/g3.113.008516)

40. Schlamp, F., Delbare, S. Y. N., Early, A. M., Wells, M. T., Basu, S., Clark, A. G. 2020 Dense time-course gene expression profiling of the Drosophila melanogaster innate immune response. bioRxiv, 2020–2026. (10.1101/2020.06.25.172452)

41. Povey, S., Cotter, S. C., Simpson, S. J., Wilson, K. 2014 Dynamics of macronutrient self-medication and illness-induced anorexia in virally infected insects. J. Anim. Ecol. 83, 245–255. (10.1111/1365-2656.12127)

42. Pham, L. N., Dionne, M. S., Shirasu-Hiza, M., Schneider, D. S. 2007 A specific primed immune response in drosophila is dependent on phagocytes. Plos Pathog 3, e26. (10.1371/journal.ppat.0030026)

43. Hilker, M., Schwachtje, J., Baier, M., Balazadeh, S., Bäurle, I., Geiselhardt, S., Hincha, D. K., Kunze, R., Mueller-Roeber, B., Rillig, M. C., Rolff, J., Romeis, T., Schmülling, T., Steppuhn, A., van Dongen, J., Whitcomb, S. J., Wurst, S., Zuther, E., Kopka, J. 2016 Priming and memory of stress responses in organisms lacking a nervous system. Biol. Rev. 91, 1118–1133. (10.1111/brv.12215)

44. Tate, A. T., Andolfatto, P., Demuth, J. P., Graham, A. L. 2017 The within-host dynamics of infection in trans-generationally primed flour beetles. Mol. Ecol. 26, 3794–3807. (10.1111/mec.14088)

45. Miller, C. V. L., Cotter, S. C. 2018 Resistance and tolerance: The role of nutrients on pathogen dynamics and infection outcomes in an insect host. J. Anim. Ecol. 87, 500–510. (10.1111/1365-2656.12763)

46. Chambers, M. C., Lightfield, K. L., Schneider, D. S., Vernick, K. D. 2012 How the fly balances its ability to combat different pathogens. Plos Pathog 8, e1002970. (10.1371/journal.ppat.1002970)

47. Leclerc, V., Pelte, N., Chamy, L. E., Martinelli, C., Ligoxygakis, P., Hoffmann, J. A., Reichhart, J. M. 2006 Prophenoloxidase activation is not required for survival to microbial infections in Drosophila. Embo Rep. 7, 231–235. (10.1038/sj.embor.7400592)

48. Chambers, M. C., Jacobson, E., Khalil, S., Lazzaro, B. P. 2014 Thorax Injury Lowers Resistance to Infection in Drosophila melanogaster. Infect. Immun. 82, 4380–4389. (10.1128/IAI.02415-14)

49. Hanson, M. A., Dostálová, A., Ceroni, C., Poidevin, M., Kondo, S., Lemaitre, B. 2019 Synergy and remarkable specificity of antimicrobial peptides in vivo using a systematic knockout approach. Elife 8. (10.7554/eLife.44341)

50. Santhanam, J., Raberg, L., Read, A. F., Savill, N. J. 2014 Immune-mediated competition in rodent malaria is most likely caused by induced changes in innate immune clearance of merozoites. Plos Comput. Biol. 10, e1003416. (10.1371/journal.pcbi.1003416)

51. Blanco-Melo, D., Nilsson-Payant, B. E., Liu, W., Uhl, S., Hoagland, D., Møller, R., Jordan, T. X., Oishi, K., Panis, M., Sachs, D., Wang, T. T., Schwartz, R. E., Lim, J. K., Albrecht, R. A., TenOever, B. R. 2020 Imbalanced host response to SARS-CoV-2 drives development of COVID-19. Cell 181, 1036–1045. (10.1016/j.cell.2020.04.026)

52. Graham, A. L. 2008 Ecological rules governing helminth microparasite coinfection. Proc. Natl. Acad. Sci. U.S.A.. 105, 566–570. (10.1073/pnas.0707221105)

